# fMRIPrep Lifespan: Extending A Robust Pipeline for Functional MRI Preprocessing to Developmental Neuroimaging

**DOI:** 10.1101/2025.05.14.654069

**Authors:** Mathias Goncalves, Julia Moser, Thomas J. Madison, rae McCollum, Jacob T. Lundquist, Begim Fayzullobekova, Lidia Hadera, Han H. N. Pham, Lucille A. Moore, Audrey Houghton, Greg Conan, Martin A. Styner, Dimitrios Alexopoulos, Christopher D. Smyser, Sally M. Stoyell, Sanju Koirala, Steven M. Nelson, Kimberly B. Weldon, Erik Lee, Robert J. M. Hermosillo, Luca Vizioli, Essa Yacoub, Gaurav H. Patel, Juan Sanchez, Kenneth Wengler, Taylor Salo, Theodore D. Satterthwaite, Jed T. Elison, Christopher J. Markiewicz, Russell A. Poldrack, Eric Feczko, Oscar Esteban, Damien A. Fair

## Abstract

The adoption of a standardized preprocessing workflow is vital for fostering community, sharing, and reproducibility. fMRIPrep has been a critical advancement towards this end, however, it is limited in its capacity to be applied to data across the lifespan, starting from infancy. Here, we introduce fMRIPrep Lifespan, an extension of fMRIPrep that extends the standardized processing from childhood to senescence to include neonatal, infant, and toddler structural and functional MRI data preprocessing. This effort involves a NiPreps integration of 1) a workflow akin to fMRIPrep optimized for MRI data in the first years of life (previously NiBabies) and 2) upstream enhancements to the entire NiPreps suite, including multi-echo data processing, modularization of workflow components, and convergence of processing with other popular workflows (ABCD-BIDS, Human Connectome Project Pipelines). Using data from the Baby Connectome Project (participants 1-43 months of age), we demonstrate that fMRIPrep Lifespan produces high-quality outputs across a wide age range. Moving forward, the scalable, modular infrastructure of fMRIPrep Lifespan will ensure adaptability to data from birth to old age while maintaining robust and reproducible frameworks for functional MRI research across the lifespan.

## Introduction

Over the past decade, functional magnetic resonance imaging (fMRI) research has developed numerous methods for data preprocessing to address research questions effectively. While these advancements have driven considerable progress, the wide variety of preprocessing options presents a challenge to reproducibility of research findings ^1,2^. Many preprocessing solutions are developed ad hoc by individual labs and often lack adherence to best practices in software engineering, including proper testing strategies, version control, and transparency, which can lead to irreproducible results ^3,4^. In addition, adopting a standardized processing pipeline facilitates communication among researchers in the field, as variation in workflows makes comparison of studies challenging, if not impossible ^1^. In response to the clear need for reproducible and standardized procedures ^4,5^, fMRIPrep was introduced as a robust preprocessing pipeline for fMRI data ^6^. To maximize reusability and shareability, fMRIPrep produces derivatives following Brain Imaging Data Structure (BIDS). fMRIPrep was quickly adopted by the field, with in-house telemetry indicating its execution over 10,000 times a week, ∼70% of which concluded with successful runs.

Ideally, fMRIPrep is generalizable across the human lifespan. However, compared to adults and aging populations, infant MRI faces a multitude of methodological challenges that need to be considered for any data processing pipeline to be compatible across the lifespan ^7,8^. For instance, neurodevelopmental changes in brain tissue myelination within the first year of life cause contrast inversion of structural MRI images and specific signal inhomogeneities ^7,9,10^ which challenge tissue segmentation and surface projection. Developmental changes in brain size and folding challenge normalization to a common space. Currently existing pipelines for adult and aging populations can therefore not simply be used for infants and young children. Data processing strategies for infants require age group-specific adaptations to succeed, and it is important to not treat infant brains as “smaller-sized adult brains”. At the same time, fMRI research in developmental populations is similarly challenged by methodological variability caused by fragmented know-how circumscribed within many labs, associated hurdles to sharing data and knowledge, and idiosyncrasies of infant imaging. Driven by large-scale studies like the developing Human Connectome Project (dHCP) ^11^, the Baby Connectome Project (BCP) ^12^, and most recently the longitudinal (birth to 10y) Healthy Brain And Child Development (HBCD) Study ^13^, fMRI data acquisition during the first 1000 days of life has quickly evolved, creating a need for standardized and reproducible data processing workflows for this age group that can extend into older ages. fMRIPrep’s already successful modular workflow ^6,14^ is ideal for tackling the challenges that come with infant fMRI data processing and, with that, extending fMRIPrep to a preprocessing workflow for the whole lifespan. While fMRIprep is able to accommodate most ages, infancy remains a major gap for standardized fMRI data processing. Therefore the first iteration of fMRPrep Lifespan focuses on optimizing data processing from birth to toddlerhood. Current and future integrations are summarized in Figure 1.

**Figure 1.**
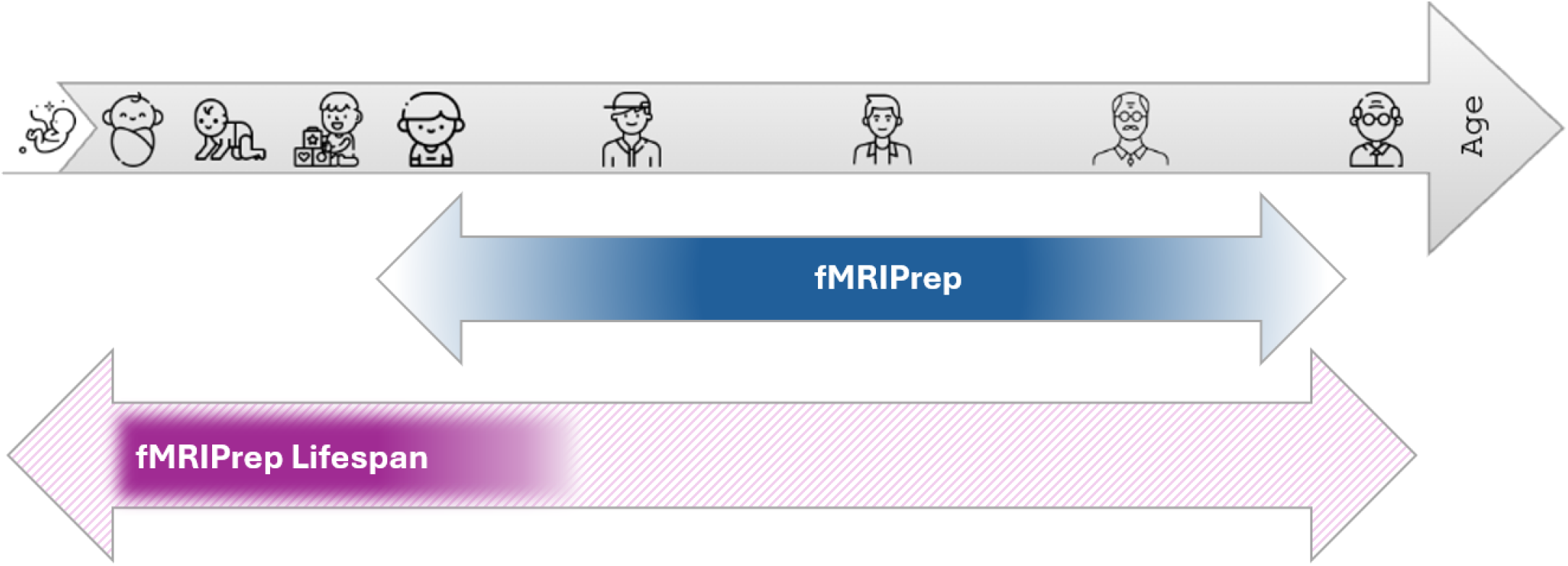
fMRIPrep Lifespan expands processing range of fMRIPrep. Solid colors represent the age range for which processing is currently optimized, color gradient shows age ranges for which processing is usable. The first version of fMRIPrep lifespan is optimized for early development after birth. Future iterations will include the whole lifespan (pattern).

The fMRIprep Lifespan pipeline integrates knowledge from existing infant workflows ^15–17^, leveraging age-specific templates and integrating alternative surface reconstruction methods, to create a robust and shareable pipeline from neonatal age to toddlerhood. A reproducible workflow to process fMRI data from all ages not only greatly benefits longitudinal studies, but also facilitates potential pooling of data.

In parallel to early-life optimizations, a number of general enhancements have proven useful to be incorporated into fMRIPrep and the NiPreps ecosystem. Enhancements include 1) optimization of computational efficiency and modularity 2) accommodation for rising acquisition sequences like multi-echo fMRI (ME), which despite being a known acquisition method for years ^18^ recently gained popularity in the field, in the context of improving data reliability in precision functional imaging ^19,20^ 3) updates to derivatives inspired by processing done in the Human Connectome Project. To quantify the impact of these updates, we evaluate the similarity of the updated derivatives to the ABCD-BIDS pipeline ^21,22^, a pipeline developed for the Adolescent Brain Cognitive Development (ABCD) project in line with HCP standards ^23^ as well as to the prior fMRIPrep-LTS (Long Term Support, 20.2.x) version.

The present paper showcases innovative developments that extend the fMRIPrep framework from a pipeline optimized for standardized data inputs from adult brains to fMRIPrep Lifespan, a versatile toolkit for fMRI preprocessing in infants, children and adults. The optimizations described in this paper significantly expand the original scope of fMRIPrep, establishing it as the first reproducible pipeline for fMRI data processing across the human lifespan.

## Results

### Age specific adaptations for infant data processing with fMRIPrep Lifespan

fMRIPrep Lifespan extends fMRIPrep’s support to MRI data preprocessing to the beginnings of the lifespan. Like its adult counterpart, it is containerized and version controlled to minimize sources of variability, relies on multiple tools within the NiPreps ecosystem, and provides easy usage, documentation, and support. fMRIPrep Lifespan produces BIDS compliant derivatives, and prioritizes longitudinal data handling by generating outputs at the session-level. fMRIPrep’s flexible workflows and modular structure makes it highly adaptable to various use cases, facilitating its adaptation to infant data preprocessing. As described below, the workflow follows a fit-transform model, to facilitate the interoperability between workflow outputs and external derivatives at various stages of processing (Figure 2). By default, the pipeline continues to use FSL FAST ^24^ for brain segmentation. However, as an alternative option for improved tissue segmentation in younger ages the pipeline supports joint label fusion (JLF). This multi-atlas approach utilizes weighted voting on expert-segmented sample images of similar developmental timepoints, improving the robustness of the segmentation ^25,26^. Additionally, users can use alternative algorithms to generate segmentations or brain masks (e.g. BIBSnet ^27^) or manually edit previously computed ones, by using fMRIPrep Lifespan’s new capability to utilize precomputed derivatives (self-made or external).

**Figure 2.**
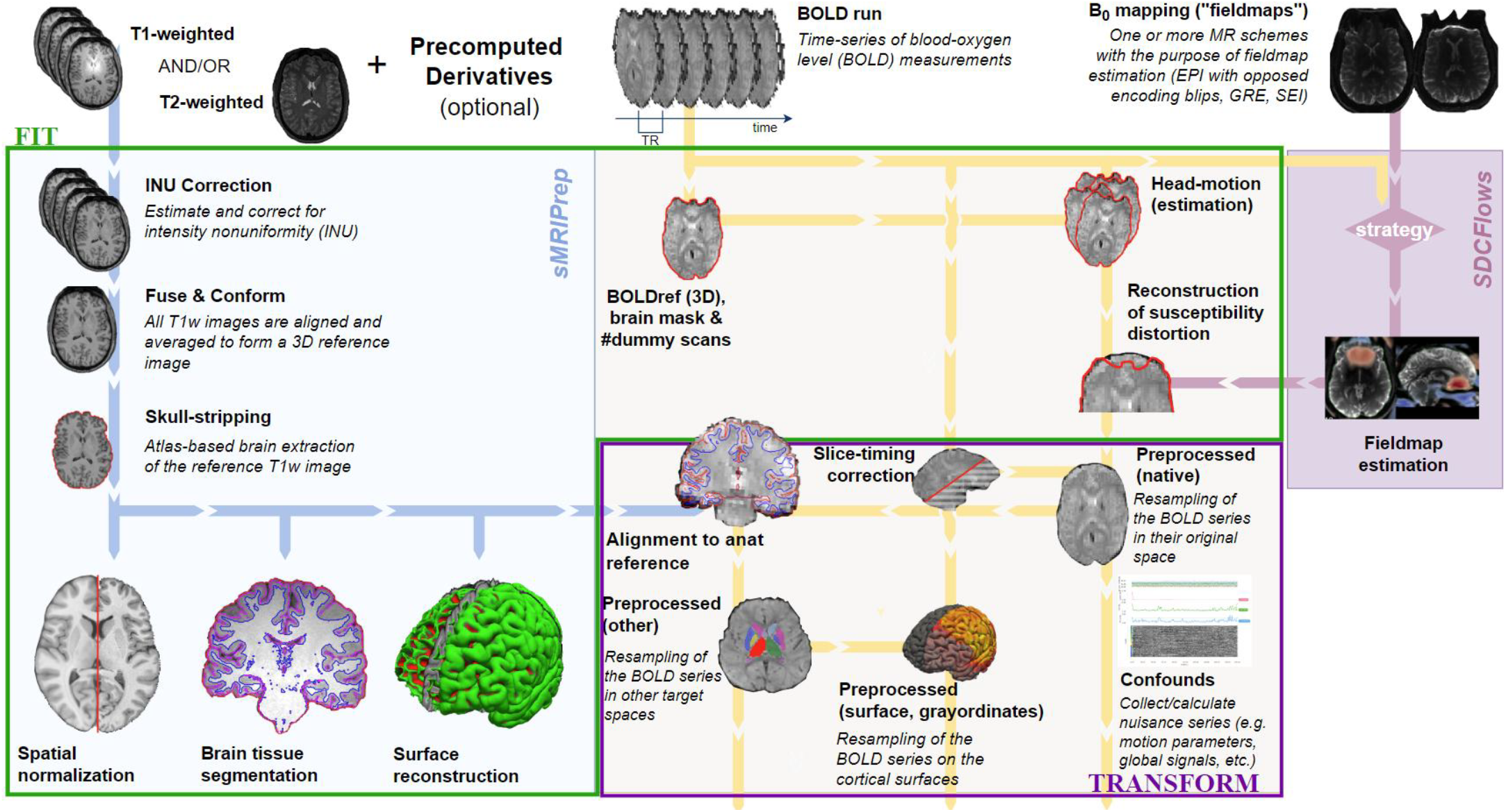
workflow schematic of fMRIPrep Lifespan showing the anatomical and functional workflow of the pipeline.

Key modifications to optimize the workflows for younger ages include the use of age-specific templates and surface reconstruction methods. Age-specific templates ^28^ are used to effectively register infant images to an intermediate, age-specific common space, which is then mapped to a standard adult stereotactic space. Treating multiple sessions of the same subject independently ensures that best-match templates are used for each scan session.

Furthermore, multiple surface reconstruction methods optimized for different age groups are available. An “auto” option is provided, which will select the surface reconstruction method best suitable based on the available data and participant age. M-CRIB-S ^29^ which operates based on T2 weighted image contrasts, is a surface reconstruction method optimized for very young infants (up to 48 weeks postmenstrual age (PMA) or 2 months) and the default method for the 0-3 months age range. This method expands fMRIPrep Lifespan to process data in cases where only a T2w image is provided. T2w images are preferred with neonatal populations (e.g. Nielsen et al. ^30^) as T1 weighted images usually provide less contrast in this early phase of life ^7^. M-CRIB-S surface reconstruction requires the user to provide a pre-computed segmentation due to the high precision required for successful surface reconstruction. The T1w is used for surface reconstruction at later time points: Infant Freesurfer ^31^ is the default for infants between 4 months and 2 years while Freesurfer recon-all ^32,33^ is implemented for infants 2 years and older. The flexible usage of either a T2w, T1w, image or both helps mitigate age-related challenges with data processing. This adaptability is particularly useful for pediatric studies where increased risk of early scanner session termination can lead to the acquisition of only a single anatomical image.

### fMRIPrep Lifespan Generates High-Quality Preprocessed Outputs

We preprocessed data from 30 randomly selected BCP study subjects, ranging in age from 1-43 months, through fMRIPrep Lifespan and assessed the quality of outputs based on manual review of surface reconstruction, spatial normalization, bias field correction, and functional alignment. Subjects were selected based on the presence of both T1w and T2w anatomical images and at least one resting state BOLD acquisition that passed raw data quality control (see Online Methods for details). For all participants, segmentations and masks were generated with BIBSnet ^27^. These externally derived derivatives were used for surface reconstruction workflows with M-CRIB-S and Infant Freesurfer, but are not supported for Freesurfer’s recon-all. For quality control purposes, data from all subjects was preprocessed with all three available surface reconstruction methods. Derivatives were quality controlled by a team of trained raters who judged the quality of surface reconstruction, spatial normalization, distortion correction and functional data alignment.

With the aforementioned challenges that are inherent to infant MRI, high quality preprocessing results can often only be achieved with manual interventions or, for some datasets, are not achievable at all. As a fully automated pipeline, fMRIPrep Lifespan produced high-quality anatomical and functional outputs for infants across the tested age range (Figure 3). The pipeline successfully completed for all 30 participants with at least one of the three surface reconstruction methods (Table 1) with processing errors being consistent with age group specific recommendations. Using the fully automated combination of BIBSNet and fMRIPrep Lifespan was successful for all participants with the Infant Freesurfer surface reconstruction. Across participants three months and younger, outputs using M-CRIB-S surface reconstruction surpassed the ones with Infant Freesurfer in quality (see Suppl. Figure 1 for an example). For most infants older than ten months, Freesurfer recon-all outputs showed accurate gray and white matter delineations but struggled with brain mask definition either missing brain regions or a too tight fit of the pial surface, cutting off parts of the gray matter (see Suppl. Figure 2 for an example). This highlights the advantage of combining a surface reconstruction method with an infant specific workflow for generating masks and segmentations, which is not possible with the present version of Freesurfer recon-all. The team of independent trained raters chose M-CRIB-S as the preferred surface reconstruction method for 4 subjects (all <= 3 months), Freesurfer recon-all for two subjects (15 and 25 months) and Infant Freesurfer for the remaining 24. Raters agreed on all but 6 participants, which were then further discussed in the team. Overall, this highlights that the surface reconstruction methods are largely optimized for the default age ranges specified by fMRIPrep Lifespan, but there are a variety of possibilities to obtain high quality outputs using fMRIPrep Lifespan’s flexible workflows. fMRIPrep Lifespan outputs from the chosen surface reconstruction methods were then further assessed for their success in spatial normalization, bias field correction and functional alignment (Figure 4; Supplementary Table 1). Most outputs were rated as either excellent or acceptable quality. Two participants’ bias field corrections were rated as poor, which was caused by significant head movement between fieldmap pair collection (Suppl.Figure 3 for an example) Head motion between field maps and runs was noted as one of the general causes of non-ideal outcomes. Even though all included datasets were rated as usable based on a raw data quality control, differences in data quality, especially between BOLD runs, was responsible for some of the variation in quality scores reported here.

**Table 1:** Number of participants (1-43 months) which successfully completed fully automated fMRIPrep Lifespan preprocessing workflow for each of the implemented surface reconstruction methods. Numbers in parentheses reflect datasets which passed quality control for surface reconstruction. Note: Externally generated segmentations (BIBSNet) were used for M-CRIBS and Infant Freesurfer but can not be used for Freesurfer recon-all. The surfaces rated as usable align well with the recommended age range for each surface reconstruction method.

|  | Completed processing | Usable surfaces | Age group for usable surfaces | Usable surfaces within default age bin | QC'ed as best surface |
| --- | --- | --- | --- | --- | --- |
| M-CRIB-S | 27 out of 30 | 6 out of 27 | $\leq 5$ months | 4 out of 4 | 4 |
| Infant Freesurfer | 30 out of 30 | 28 out of 30 | $\geq 3$ months | 16 out of 16 | 24 |
| Freesurfer recon-all | 22 out of 30 | 14 out of 22 | $\geq 10$ months | 7 out of 10 | 2 |

**Figure 3.**
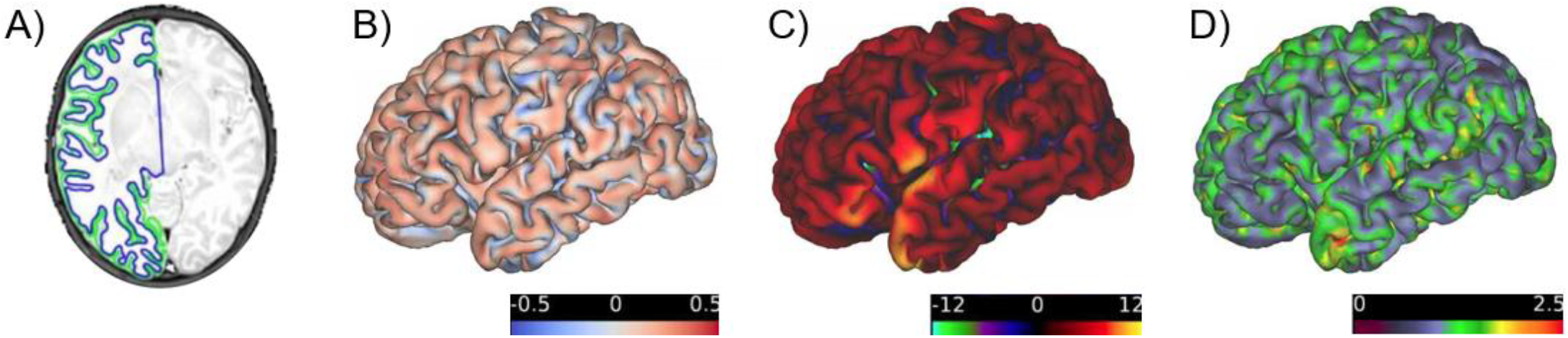
Example fMRIPrep Lifespan outputs in a 1 month old. A) T2w image with outlines of white matter (blue) and pial surface (green) B) Curvature map C) Sulcal depth map D) Cortical thickness map.

**Figure 4.**
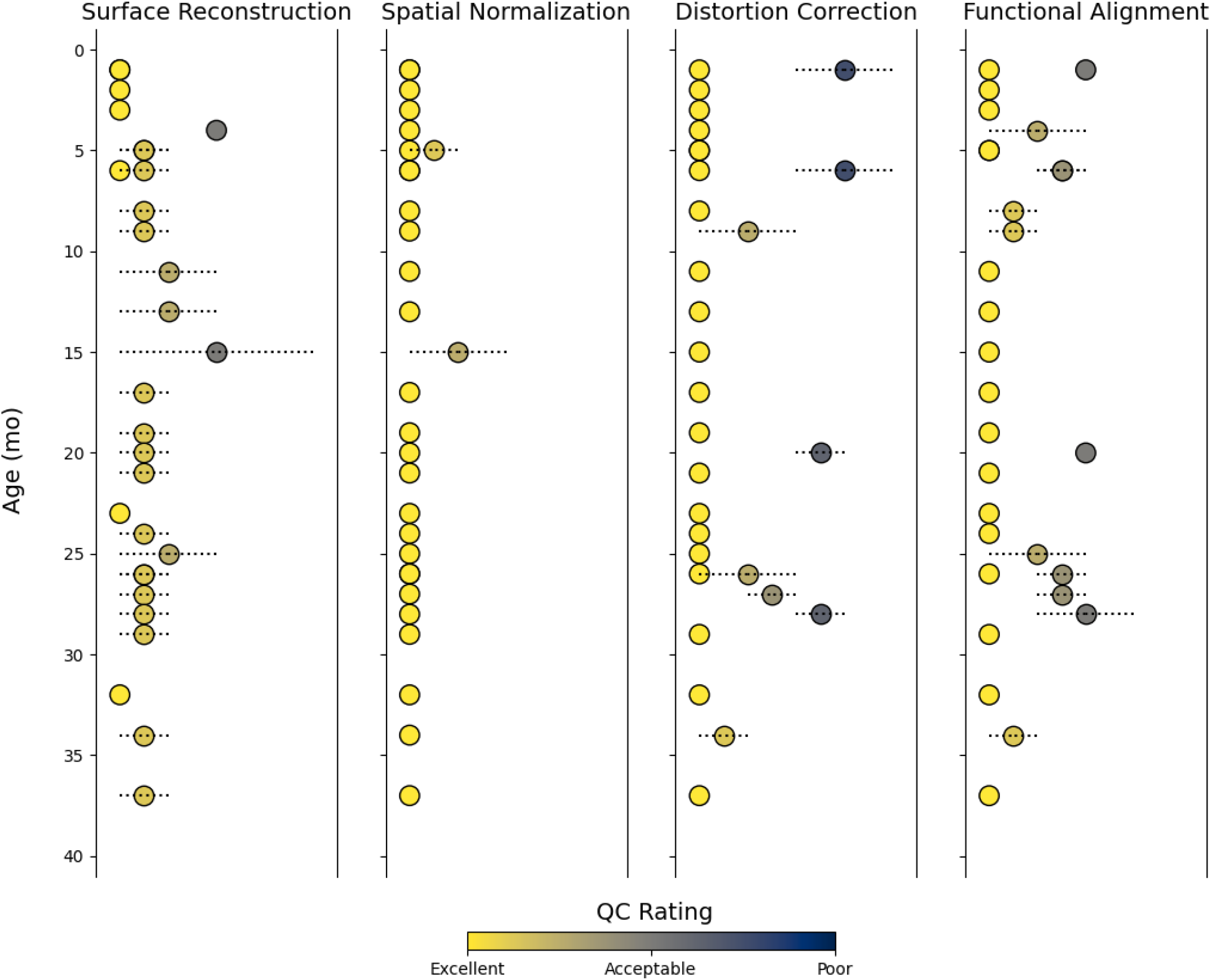
Quality control outcomes for all participants by age. Results were generated by six trained raters, each output was reviewed by two independent raters. Ratings were performed on a scale from 1-3, differentiating between images with high quality, usable with room for improvement and not of acceptable quality. Dots containing a line show discrepancy between reviewer ratings.

As a preliminary test of fMRIPrep Lifespan’s ability to operate across a wider range of ages, we explored the option of processing a subset of ABCD subjects (N=15, ages 9-11 years) with fMRIPrep Lifespan, using Freesurfer recon-all for surface reconstruction. Despite the lack of optimization of the current version of fMRIPrep Lifespan for this age group, 12/15 test cases finished successfully, albeit with varying output quality. Failures were based on problems aligning data with the non-age matched anatomical templates. Future releases of fMRIPrep Lifespan will include matching templates for children in this age range.

### Age agnostic updates enhancing the entire NiPreps suite

In addition to infant-specific changes, several age agnostic enhancements have been implemented in the NiPreps ecosystem as a whole. Notably, the fMRIPrep-23.2.X series includes a variety of changes, encompassing from the restructuring of the workflow, changes to the processing of each modality, to alignment of outputs with commonly used standards (see Online Methods and Supplementary Table 2 for details). Enhancements to data preprocessing include support for multi-echo data inputs, a reworked susceptibility distortion correction workflow, the option to mask out high-variance voxels, and improvements to data surface mapping. Each of these improvements are described in detail below:

#### Workflow structure

The revised workflow architecture represents a fit-transform model, composed of two sub-workflows: the “fit” stage involves generating computationally expensive outputs, such as registrations and surface reconstruction, which are subsequently applied on the raw data during the “transform” stage. Splitting the workflow into these two stages distributes tasks to increase computational efficiency, decreasing fMRIPrep’s runtime up to 98% ^34^ while also increasing accessibility to researchers with less computational resources. Additionally, this overhauled workflow architecture exposes ingestion points through the workflow, allowing for both reuse of fMRIPrep-made derivatives, as well as externally computed derivatives into the pipeline.

#### Multi-echo support

fMRIPrep now automatically detects multi-echo data in BIDS input data and adjusts its workflows accordingly. fMRIPrep uses the first echo when computing spatial transformations, as it tends to have the least signal dropout, and applies these to all echos. Following slice-timing, head motion, and susceptibility distortion correction, an “optimal combination” of echos is generated using Tedana ^35^, and resampled into any desired target spaces. Alternatively, users can specify for the minimally processed individual echos to be output, allowing for echo-specific analyses and/or further denoising using Tedana or other multi-echo tooling. Maps of T2* relaxation times that can be overlaid with the BOLD data are generated in addition to standard derivatives (Suppl. Figure 4).

#### Susceptibility distortion correction (SDC)

SDCFlows, the NiPreps utility used by fMRIPrep for susceptibility distortion estimation and correction, has been overhauled with a new uniform Application Programming Interface (API) for all supported fieldmap estimation methods, facilitating comparison and combinations of distortion correction strategies. SDCFlows provides a diverse range of estimation methods, with the best fit selected based on the data available. These methods include direct B_0_ mapping, phase difference B_0_ estimation, PEPOLAR (Phase Encoding POLARity), and fieldmap-less SDC. In addition to the previous implementation using AFNI’s 3dQwarp, a new workflow utilizing FSL’s “topup” ^36^ is available to estimate with two or more differing phase encoding directions (Suppl. Figure 5). Following the fit-transform paradigm, the fit stage of SDCFlows estimates a B_0_ field inhomogeneity map in Hertz -generally referred to as “fieldmap” - that is then regularized with a B-Spline basis grid. Anatomical references are then generated to project the fieldmap into target BOLD space for correction. The transform stage of SDCFlows implements the fit step for the respective field map estimation method. SDCFlows includes workflows for the correction of distorted BOLD scans that co-register the fieldmap’s anatomical reference to the target, interpolates the fieldmap on the grid of the target from the B-Spline coefficients through the co-registration transform, and finally resamples the BOLD scan after converting the fieldmap from Hertz into dense displacements field. SDCFlows also implements a nonlinear spatial transform that can be combined with other transformations to minimize data resamplings with *NiTransforms* ^37^.

#### Improved volume-to-surface mappings and grayordinates (mixed volume and surface data representations)

To improve similarity to HCP workflow outputs as well as BOLD data registration, fMRIPrep now generates anatomical derivatives (white matter, sulcal depth, curvature, etc) in “grayordinates” ^23^ that refer to the fsLR template ^38^. An additional workflow has been integrated to generate an anatomical cortical ribbon volume based on the signed distance function of the white and pial surfaces. Furthermore, direct resampling from the subject’s FreeSurfer native mesh (“fsnative”) to fsLR is now implemented, and a Connectome Workbench workflow using subject-specific cortical ribbons produces more accurate surfaces. (Suppl. Figure 6). The surface mapping is additionally enhanced with Multimodal Surface Matching (MSM); ^39,40^), also in line with HCP workflows ^41^. Here, MSM optimizes the registration of the subject’s native surface sphere to a standard template reference by improving the overlap of sulcal geometry between subject and reference and reducing distortion of edge lengths, triangle areas, and triangle shapes. An optional workflow is added to exclude voxels with local peaks of temporal variation from the BOLD time-series (“--project-goodvoxels”; Suppl. Figure 7). Voxels with local peaks of temporal variation are often near the edge of the brain parenchyma or contain large blood vessels and removing them eliminates a gyral bias in high coefficient of variation ^23^. Together, these additions facilitate comparison of data derivatives with those from HCP-style pipelines ^23,41^ while maintaining compliance with the BIDS specification for imaging data derivatives (Figure 1).

#### Improved downstream compatibility with XCP-D for denoising

The updated fMRIPrep furthermore generates outputs suitable for ingress by XCP-D ^42^, a highly configurable pipeline for functional connectivity post-processing. XCP-D performs additional data cleaning steps (e.g. respiratory filter, global signal regression) and provides options to output denoised data in standard volume spaces, CIFTI dense grayordinates, or parcellated grayordinates for various parcellation schemes.

### Enhancements lead to improved convergence between fMRIPrep and ABCD-BIDS

We compared fMRI outputs in the form of a parcellated correlation matrix between the enhanced version (fMRIPrep-23.2.1), the LTS version (fMRIPrep-20.2.7) and ABCD-BIDS using 50 randomly selected participants from the ABCD study (see Online Methods for details). Similarity of outputs was quantified based on intraclass correlation coefficient (ICC) and Pearson correlation (corr) between matrices (Figure 5). We found that fMRIPrep-23.2.1 outputs were more similar to ABCD-BIDS (mean ICC = 0.93, SD = 0.11; mean corr = 0.92, SD = 0.03) than fMRIprep-LTS to ABCD-BIDS (mean ICC = 0.88, SD = 0.08; mean corr = 0.81, SD = 0.05), displaying a significant increase in similarity in both ICC (p<0.001) and matrix correlation (p<0.001). Similarity between outputs from fMRIprep-23.2.1 and ABCD-BIDS surpass the similarity between fMRIprep-23.2.1 and fMRIPrep LTS derivatives (mean ICC = 0.86, SD = 0.12; mean corr = 0.84, SD = 0.04).

**Figure 5.**
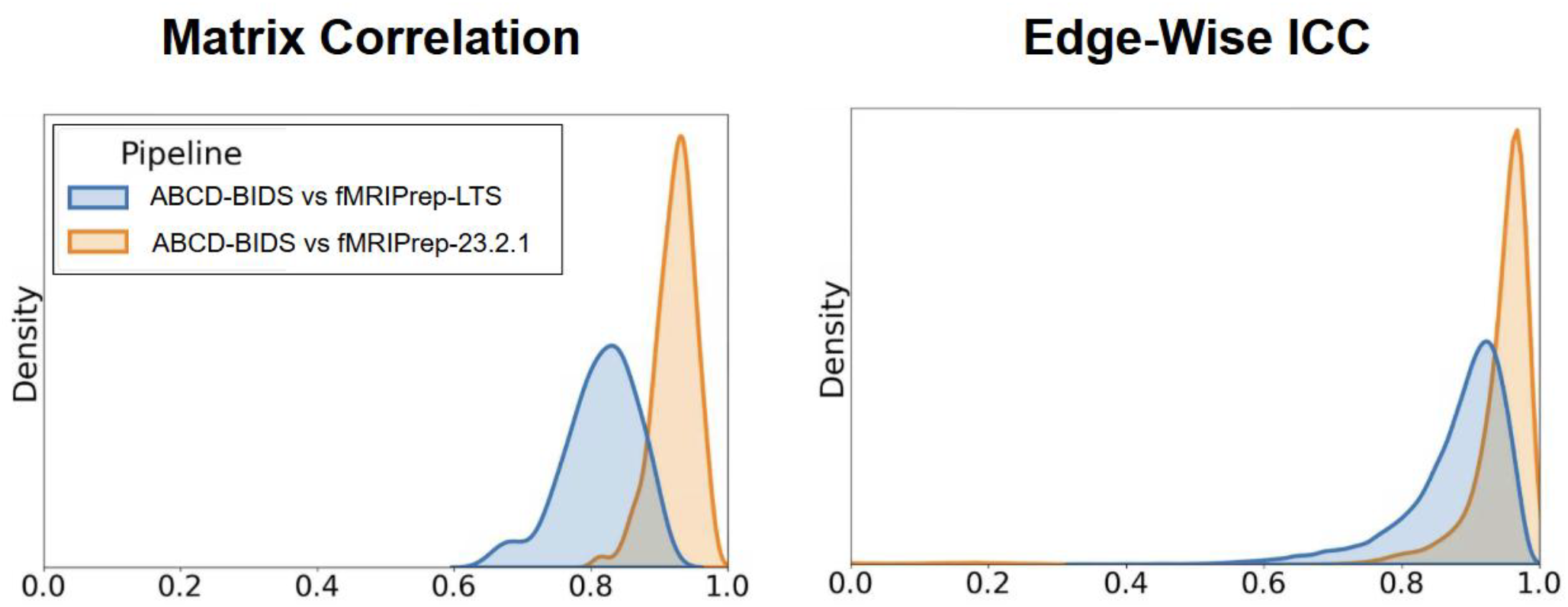
Comparison of different versions of fMRIPrep to ABCD-BIDS. fMRIPrep-23.2.1 is the recent version discussed in this paper, LTS is (20.2.7) the most recent of the long term support version from 2020. Metrics are Pearson correlation and ICC, calculated on parcellated data of 50 ABCD subjects. The comparison for each metric is specified on the left.

## Discussion

In this paper we introduce fMRIPrep Lifespan, which includes a workflow that is optimized for MRI data of neonates, infants and toddlers. Using a random set of test datasets from the BCP (participants scanned between 1-43 months postpartum) we showed that fMRIPrep Lifespan produces high-quality outputs across a wide age range. fMRIPrep Lifespan includes enhancements to the entire NiPreps suite (for infants and adults), including multi-echo data processing, modularization of workflow components, and convergence of processing with other popular workflows. A comparison performed in this paper shows that the enhanced adult fMRIPrep version produces derivatives similar to ABCD-BIDS, which is in line with standards established during the Human Connectome Project.

fMRIPrep Lifespan is an important extension that replicates fMRIPrep’s approach and facilitates infant data processing by simplifying usability and increasing speed while addressing the distinctive challenges posed by infant data. This is a particularly timely adaptation in light of the recent increase in infant neuroimaging research and the start of the large-scale HBCD study ^13^. The workflow is adapted to the needs of infant imaging, particularly with its different structural workflow which includes age specific templates and the option to process data with either a T1w, a T2w or both anatomical images and the integration of three different surface reconstruction methods. This level of adaptiveness is key given the rapid rate of brain development in the first year of life ^7,43^. It provides flexibility to feed custom inputs for segmentations and masks as expert intervention is often required to manually correct segmentations if brain tissue is not properly delineated by automated algorithms. In comparison to other workflows developed for infant imaging ^15–17^, it is open source and shareable, covers a wide age range (not only the neonatal period) and its modular architecture facilitates continuous improvements.

fMRIPrep Lifespan produces high quality outputs across a wide age range as demonstrated in our use case example with BCP participants 1-43 months of age. All participants could be successfully processed, passing quality control, with at least one of the three surface reconstruction methods. Suboptimal results with using a surface reconstruction method that is not recommended for a given age group (e.g. M-CRIB-S for infants past 3 months) highlight the necessity of having an age specific flexible workflow. M-CRIB-S is optimized for the neonatal period and assumes a T2w image with a contrast that is inverted compared to images from older children or adults as an input. This will limit its chances for successful surface reconstruction in infants past 3 months, who start to have changes in the T2w image contrast. On the other hand, Freesurfer’s recon-all is optimized for adult-like T1w images which will limit its utility for infants in or before the isointense period. In our test case, Infant Freesurfer showed the highest level of success across the board. The main driver for that is the use of high quality externally generated segmentations (in comparison to the Freesurfer recon-all workflow) in combination with Freesurfer having less age specific priors (in comparison to M-CRIB-S). The Freesurfer recon-all workflow showed benefits in detailed white matter delineations in many subjects with more adult like T1w images (>12 months) and an enhancement for future versions of fMRIPrep Lifespan could therefore include a combination of Freesurfer recon-all surface reconstruction with the ingestion of externally generated masks and segmentations. All segmentations used within our workflow were generated fully automated. Further manual corrections to refine segmentations can be used to optimize surface reconstruction results.

The fMRIPrep Lifespan testing approach used in this paper used real world data for pipeline testing. Even though all included participants passed a raw data quality control, for some subjects optimal results could still not be achieved. Reproducible pipelines enable researchers to investigate the relationship between data quality, pipeline options, and processing output quality in order to further refine pipelines as well as refine data acquisition protocols and processes. Based on the observation of problems with the quality of distortion correction caused by suboptimal fieldmap pairs one promising future pipeline refinement will be the improvement of synthetic bias field correction for infant datasets.The results presented in this paper demonstrate the benefit of the flexible workflows implemented in fMRIPrep Lifespan, which allows it to cater to the needs of different age groups or available high quality images.

The extensive improvements made to fMRIPrep not only produce higher quality outputs, through additions such as sulcus-based surface matching ^39^ and optional “goodvoxels” masking ^23,41^ to improve fMRI surface mapping, but also improve shareability of data by complying with BIDS standards, and comparability with HCP-style outputs by including HCP-style processing methods and output formats. HCP-style pipeline outputs ^23,41^ are widely used in the field of neuroimaging for analysis including, for example, derivatives provided for analysis in CIFTI grayordinate standard spaces. The support of automated handling of ME-fMRI data inputs serves a recent rise in popularity of this method (e.g. Lynch et al. ^19^) and speaks to fMRIPrep’s increased versatility. The increase in options within the fMRIPrep framework allows the use of the same pipeline across a wide range of studies and thereby contributes to reproducible research and facilitates comparability and pooling of data derivatives. With the changes described here, fMRIPrep produces results that are very similar to those generated with ABCD-BIDS while being a robust and shareable pipeline which is at the same time modular and adaptable. This will allow fMRIPrep to grow with the needs of the field. Furthermore, fMRIPrep is well documented and supported and can therefore be easily adapted by neuroimaging researchers from different backgrounds. Alignment with standards for software documentation, infrastructure, and testing ^4^ via the NMIND checklist tool (nmind.org/proceedings) helps to maintain this standard and avoid common pitfalls for reproducibility ^3^. fMRIPreps compatibility with post-processing options such as XCP-D ^42^ allows to integrate it into a completely reproducible workflow for resting state functional connectivity data which can be used across the whole lifespan.

Community adoption is indicative of there being a real need for a streamlined and robust minimal preprocessing pipeline for MRI data across the lifespan. However, as shown with neonates and infants, alternative techniques and optimizations are required to best serve the specifics of each data. The formation and ongoing development of fMRIPrep Lifespan, in parallel with other population-specific fMRIPrep forks (fMRIPrep-rodents ^44^), make evident a growing need for a plugin or extension-based approach. This effort has already begun with the modularization of independent modality processing (sMRIPrep for structural, SDCFlows for susceptibility distortion correction), but further infrastructure is needed to provide fine-grain connection points between internal workflows. Incorporating this extension-based approach will greatly reduce the developmental burden of porting upstream changes across extension pipelines. Our preliminary test with 15 ABCD subjects shows that the extension of fMRIPrep Lifespan to further age groups is not a far reach. In the framework of NiPreps, this reproducible workflow for all ages can be extended to other modalities such as diffusion MRI or MRI data quality control ^14^.

With ongoing development, the pipeline will be further optimized to be specific for all age groups, by integrating fitting templates for older children, older adults and potentially fetal MRI. Moving forward, the framework will be extended based on the needs in the field, ideally through continuous user feedback and support. One example is the implementation of MSMSulc in the fMRIPrep Lifespan workflow and the ability to use the synthetic field map feature of SDCFlows. A future direction, addressing growing needs in the field, will be an optimization of fMRIPrep for fMRI data acquired at 7 Tesla. Ultra-high field fMRI is gaining in popularity and only recently its use has been expanded to infant populations ^45^. With its modular structure, which allows for easy implementation of new developments, and its current workflows for supporting developmental fMRI research starting in newborns, fMRIPrep Lifespan is the optimal tool to provide the neuroimaging community with a reproducible one-stop tool for fMRI preprocessing.

## Online Methods

### fMRIPrep Lifespan

fMRIPrep Lifespan is an expansion of fMRIPrep ^6,46^, with the goal of providing preprocessing for the whole human lifespan. Currently, it is a separate pipeline optimized for infant fMRI data processing. The underlying workflow engine used in fMRIPrep Lifespan is Nipype 1.8.6 ^47^ and many internal operations of fMRIPrep Lifespan use Nilearn 0.10.4 ^48^. The input dataset is processed by subject and session groupings to ensure closest-match templates are used for each time-point. It can be subdivided into an anatomical and a functional workflow as detailed blow:

#### Anatomical workflow

Both a T1-weighted and T2-weighted image are recommended for best performance, but the workflow only requires a single anatomical image. For each anatomical available, a canonically-oriented, structural reference is created. If multiple runs are found within a session, the intensity-nonuniformity-corrected versions are fused into a reference map of the subject with mri_robust_template. If multiple anatomicals are found, the anatomical images are coregistered, and then the primary reference image used will be determined based on data availability, participant age, and desired surface reconstruction method.

The anatomical reference is denoised (ANTs DenoiseImage), corrected for intensity nonuniformity (ANTs N4BiasFieldCorrection; see e.g. ^49^) and undergoes a modified antsBrainExtraction workflow where it is co-registered to an age-matched template (two options available: the UNCInfant 0-1-2 template (default) ^50^ and the MNIInfant (0-4.5 yr) template ^28^). If provided, the workflow can use a pre-computed brain mask throughout the rest of the workflow.

If no external segmentation is provided, brain tissues - cerebrospinal fluid (CSF), white matter (WM), and gray matter (GM) - are segmented from the reference, brain-extracted anatomical image using Joint Label Fusion (if age-matched atlases are available) ^25^. Otherwise, FSL FAST is used ^51^ (As contrasts vary, this is not ideal.)

There are three surface reconstruction methods available in fMRIPrep Lifespan, each intended to maximize performance based on brain morphometry during an age range. *M-CRIB-S* ^29^. MCRIBReconAll is intended for cases where T1w tissue contrast is poor, likely due to age (0-3 months), and uses the T2w reference image. *FreeSurfer* ^*32*^. tools’ infant_recon_all (4-24 months; within *Infant FreeSurfer* ^31^ and recon-all (24 months and onward; standard *FreeSurfer*) use the T1w reference to generate surfaces.

Volume-based data is spatially normalized into infant standard space MNIInfant (0-4.5 yr) template ^28^) through nonlinear registration with antsRegistration (ANTs), using brain-extracted versions of both T1w (or T2w) reference and the T1w (or T2w) template. The templates are accessed with TemplateFlow ^52^. Other spaces can be requested using the ‘output spaces’ option.

#### Functional workflow

The first step in fMRIPrep Lifespan’s functional workflow is the generation of a reference volume (17th volume as in the DCAN labs infant-abcd-bids pipeline ^16^) for use in head motion correction. Head-motion parameters with respect to the BOLD reference (transformation matrices, and six corresponding rotation and translation parameters) are estimated before any spatiotemporal filtering using MCFLIRT (FSL, ^53^).

##### SDCFlows

In the present version of fMRIPrep Lifespan, the fieldmap or B0-nonuniformity map can be estimated based on three different strategies in the “fit” stage of SDC Flows: *1) Direct B0 mapping:* for example, with Spiral-Echo Imaging which reconstruct an estimate of the fieldmap in Hz *2) Phase-difference B0 estimation:* calculated from two subsequent Gradient-Recalled Echoes ^54 &^ *3) PEPolar estimation strategy:* with two (or more) opposing echo-planar imaging (EPI) references using FSL’s topup ^36^. In the “transform” stage, an estimated fieldmap is then aligned with rigid-registration (ANTs) to the target EPI (echo-planar imaging) reference EPI. The field coefficients are mapped on to the reference EPI using this transform. The BOLD reference is co-registered to the anatomical reference using bbregister (FreeSurfer) which implements boundary-based registration ^55^. Co-registration is configured with six degrees of freedom. The aligned T2w image is used for initial co-registration (unless T2w image is unavailable, then the T1w image is used).

##### Calculation of confounds

Several confounding time-series are calculated based on the preprocessed BOLD: framewise displacement (FD), DVARS (spatial root mean square of the data after temporal differencing) and three region-wise global signals. FD is computed using two formulations following Power (absolute sum of relative motions, ^56^) and Jenkinson (relative root mean square displacement between affines, ^57^). FD and DVARS are calculated for each functional run, both using their implementations in Nipype (following the definitions by ^56^). The three global signals are extracted within the CSF, WM, and the whole-brain masks. Additionally, a set of physiological regressors are extracted to allow for component-based noise correction (CompCor, ^58^).

Principal components are estimated after high-pass filtering the preprocessed BOLD time-series (using a discrete cosine filter with 128s cut-off) for the two CompCor variants: temporal (tCompCor) and anatomical (aCompCor). tCompCor components are then calculated from the top 2% variable voxels within the brain mask. For aCompCor, three probabilistic masks (CSF, WM and combined CSF+WM) are generated in anatomical space. The implementation differs from that of Behzadi et al. in that instead of eroding the masks by 2 pixels on BOLD space, a mask of pixels that likely contain a volume fraction of GM is subtracted from the aCompCor masks. This mask is obtained by dilating a GM mask extracted from the segmentation, and it ensures components are not extracted from voxels containing a minimal fraction of GM. Finally, these masks are resampled into BOLD space and binarized by thresholding at 0.99 (as in the original implementation).

Components are also calculated separately within the WM and CSF masks. For each CompCor decomposition, the k components with the largest singular values are retained, such that the retained components’ time series are sufficient to explain 50 percent of variance across the nuisance mask (CSF, WM, combined, or temporal). The remaining components are dropped from consideration. The head-motion estimates calculated in the correction step are also placed within the corresponding confounds file. The confound time series derived from head motion estimates and global signals are expanded with the inclusion of temporal derivatives and quadratic terms for each ^59^. Frames that exceeded a threshold of 0.5 mm FD or 1.5 standardized DVARS are annotated as motion outliers. Additional nuisance timeseries are calculated by means of principal components analysis of the signal found within a thin band (crown) of voxels around the edge of the brain, as proposed by ^60^.

##### BOLD resampling

For sampling to surface spaces, BOLD time-series are registered to the subject’s primary anatomical volume template and then projected using the ribbon-constrained volume-to-surface-mapping function of Connectome Workbench, targeting the subject’s fsnative mesh ^23^. If the “--project-goodvoxels” flag is used, a “goodvoxels” mask is applied to the BOLD time-series to exclude voxels whose time-series have a locally high coefficient of variation from being projected ^23^. Data sampled from subject’s anatomical template space gets resampled onto the left/right-symmetric template fsLR using Connectome Workbench ^23^. If dense CIFTI format outputs are desired, grayordinates files ^23^ containing 91k samples are generated, with registration from fsnative to fsLR 32k template using MSM.

Volume-space resamplings can be performed with a single interpolation step by composing all the pertinent transformations (i.e. head-motion transform matrices, susceptibility distortion correction when available, and co-registrations to anatomical and output spaces). Gridded (volumetric) resamplings are performed using nitransforms ^61^, configured with cubic B-spline interpolation.

### fMRIPrep: evolution from LTS to current version

#### Restructured workflow

##### Fit-transform model

The workflow architecture is composed of two sub-workflows for each modality processing: anatomical, fieldmap, and functional. These sub-workflows, referred to as “fit” and “transform” stages, are used to conceptually separate tasks and outputs. First, in the “fit” stage, computationally expensive tasks, such as registration, segmentation, and reconstruction, can be run and produce small and thus easy to distribute outputs. Then, during the “transform” stage, these outputs, along with the raw dataset, can be consumed to produce deterministic outputs, such as the data resampled into a different space. This separation of processing can be used to spare the user from excessive use of computational resources and disk space.

#### Processing changes

Many improvements have been made to fMRIPrep since the LTS, with some of the most notable being:

##### Enhanced multi-echo support

Leveraging Tedana ^35^, fMRIPrep processes all supplied single echo time series, using the BOLD reference mask and optimally combines the data in a T2* weighted fashion. By default, a nonlinear regression is used to fit curves for estimating T2* times. Optionally, if “--me-output-echos” is requested, each echo is motion corrected, and slice-timing and susceptibility distortion corrected, if available, but time series are not optimally combined based on the T2* map. These outputs can for example be used to perform additional denoising using ME-ICA ^62^. Additionally, a new reportlet was added comparing the T2* map with the BOLD reference, superimposing the anatomical gray-matter mask on both. A histogram plot shows the distribution of values within the mask.

##### Susceptibility distortion correction

A comprehensive overhaul of SDCFlows, the Python library of NiPype-based workflows used by fMRIPrep to preprocess *B0* mapping data, estimate corresponding fieldmaps and correct for susceptibility distortions. SDCFlows now represents all map estimations with a B-Spline basis, ensuring a common basis between different correction strategies, facilitating combination and comparison of the strategies supported. Changeover to a fit-transform approach splits estimation and correction into separate, standalone processes.

Furthermore, when acquiring two or more Spin Echo EPI scans with different phase encoding directions, also known as PEPolar, the default correction strategy is now an implementation of FSL’s topup utility, replacing the prior method using AFNI’s 3dQwarp. The new TOPUP-based strategy is robust to various acquisition schemes, supporting PEPolar inputs with different effective echo spacings, total readout times, and orthogonal phase-encoding directions (none of which were supported under 3dQwarp); in addition, these changes bring fMRIPrep’s methods more in line with ABCD-BIDS and other HCP-based pipelines.

#### SyN Correction

This workflow takes a skull-stripped anatomical image and a reference EPI image, and estimates a field of nonlinear displacements that accounts for susceptibility-derived distortions. To more accurately estimate the warping on typically distorted regions, this implementation uses an average mapping ^63^. This process is divided into two steps - first, a reference map is calculated for each image to be aligned, as well as intensity clipping and deobliquing. Secondly, the images are nonlinearly registered, using ANTs’ SyN method. To aid the Mattes mutual information cost function, the registration scheme is set up in multi-channel mode, using laplacian-filtered derivatives in the second channel. The anatomical image is used as the fixed image, and therefore, the registration process estimates the transformation function from unwarped (anatomically correct) coordinates to distorted coordinates. The resulting transform is converted into a B_0_ fieldmap in Hz and stored as an “fmap/” output.

#### CIFTI outputs

To more closely resemble outputs from the Human Connectome Project Pipelines, significant additions have been added to fMRIPrep’s CIFTI workflow. First, FreeSurfer morphometric maps of cortical thickness, curvature, and sulcal folding (average convexity) are produced ^64^, and are converted to native-space GIFTI and optionally fsLR-space CIFTI formats. GIFTI files follow FreeSurfer conventions (negative curvature and negative sulc are gyral) while the fsLR-space CIFTI-2 files follow HCP conventions (positive curvature and sulcal convexity are gyral). GIFTI files are intended to be minimal conversions from FreeSurfer, while the CIFTI-2 dscalars in fsLR space are directly comparable to HCP outputs. In addition, workflows were added to 1) generate an anatomical cortical ribbon mask, based on the signed distance function of the white and pial surfaces and 2) apply this mask to the BOLD time-series to exclude voxels with local peaks of temporal variation. This option can be enabled with the “--project-goodvoxels” flag.

To improve spherical registration to fsLR surfaces, Multimodal Surface Matching (MSM) was incorporated into the workflow, leveraging the FreeSurfer sulcal maps to improve folding alignment. The implementation is based on the HCP Pipelines’ MSMSulc pipeline with a “strain-based” cost function, which has demonstrated reduced distortions in edge lengths, triangle areas, and triangle shapes compared to FreeSurfer’s spherical registration method ^40^.

## Datasets

### BCP

30 subjects from the BCP ^12^ dataset were selected to demonstrate processing outcomes. Subjects that were chosen had to pass the raw MRI visual assessment for excessive motion, insufficient coverage, and/or ghosting. Subjects were primarily selected based on whether they had at least one run of both T1w and T2w anatomical images, as well as at least one run of resting state BOLD-AP/PA data with corresponding field map runs (AP/PA). We randomly sampled 10 subjects (with the only restriction being the previously stated data availability criteria) from three age bins (0-3 months, 4-24 months, 25-44 months) to demonstrate the application of three different surface reconstruction methods supported in fMRIPrep Lifespan. The subjects’ mean age was 17.9 months, 4 were longitudinal participants (two sessions from the same participant were used in our analyses), 17 participants were identified as female (2 longitudinal), 19 were white (4 longitudinal). Data was collected at two sites: University of North Carolina (UNC), Chapel Hill and University of Minnesota (UMN). Both sites used a Siemens 3T Prisma (32-channel coil) scanner. The imaging modalities collected for the BCP study were structural, functional (resting state) and diffusion, but diffusion data cannot be processed by fMRIPrep Lifespan and was therefore not included in our analyses. The scanning sequences for the MPRAGE T1w included the following parameters: echo time (TE) = 2.24 msec, repetition time (TR) = 2400 msec, flip angle = 8, resolution = 0.8 mm^3^, multi-band factor (MB) = 1. The T2w images used a 3D variable flip angle turbo spin-echo sequence (Siemens-Space, turbo factor = 314, echo train length = 1166 msec; ^65^). The remaining parameters for the T2w modality were as follows: TE = 564 msec, TR = 3200 msec, resolution = 0.8 mm^3^, acceleration factor (AF) = 2. The scanning sequence parameters for the resting state BOLD-AP/PA modality are as follows: TE = 37 msec, TR = 800 msec, flip angle = 52, resolution = 2 mm isotropic, MB = 8. Finally, the scanning sequence parameters for the field map AP/PA modality are as follows: TE = 66 msec, TR = 8000 msec, flip angle = 90, resolution = 2 mm isotropic, MB = 1.

### ABCD

50 subjects from the ABCD dataset were randomly selected using the following criteria: all subjects passed raw data quality control, were from the baseline session and from the same collection site (site02), with data acquired using the same scanner manufacturer, model, and software version (SIEMENS Prisma Fit, syngo MR E11). The sample had a mean age of 10.2 years (8.9-11 years) or 122 months (107-132 months), with 23 females and 46 white subjects. For details on ABCD data acquisition see ^66^.

## Analysis methods

### BCP

#### Data processing

Segmentations were generated using BIBSnet ^27^ using each subject’s T1w and T2w anatomical image. These segmentations were used as external derivatives for fMRIPrep Lifespan (workflow described above). All 30 datasets were processed with all three available surface reconstruction methods.

#### Analysis

Data outputs generated by fMRIPrep Lifespan version 25.0.1 were quality controlled by two trained independent raters per dataset. The team of raters were authors JM, JTL, rMC, HHNP, BF and SMS. Surface reconstruction was evaluated using the native space T1 and T2 with the constructed pial and white matter surfaces overlayed. Raters assessed all three surface reconstruction methods for each dataset and chose the best method for a given dataset. Disagreements between independent raters were discussed within the team until a consensus was reached. Further quality assessment of derivatives was performed from outputs generated with this most fitting method. The following dimensions were evaluated using outputs from the fMRIPrep Lifespan derivatives structure: spatial normalization, bias field correction, and functional alignment. Spatial normalization and bias field correction were assessed using the HTML summary document. Functional alignment was assessed using the boldref in the T1 space alongside the pial and white matter surfaces. These two evaluations were done using Connectome Workbench for visualization. Raters used the categories “excellent” (1) to indicate overall good outputs, “acceptable” (2) to indicate usable outputs with some room for improvement, and “poor” (3) for unusable outputs. Raters additionally took notes about quality issues in inputs or outputs.

### ABCD

#### Data processing

Data were preprocessed using fMRIPrep 23.2.1, fMRIprep-LTS, and ABCD-BIDS ^22^. Preprocessed data was divided into parcells, using Workbench command ^23^ and a template of the Gordon parcellation scheme ^67^. *Analysis:* Pearson correlations were calculated in Matlab by vectorizing the upper triangles of the connectivity matrices and using the “corr” function. ICC was calculated using the PyReliMRI package ^68^. The conn_icc.py script was used to run the edgewise_icc function using icc_type 3 (see Github for reference).

## Data and code availability

Data: Both BCP and ABCD datasets are publicly available through the National Data Archive. fMRIPrep and fMRIPrep Lifespan:

fMRIPrep DockerHub, fMRIPrep GitHub, fMRIPrep readthedocs fMRIPrep lifespan (NiBabies) DockerHub, fMRIPrep lifespan (NiBabies) GitHub,fMRIPrep lifespan (NiBabies) documentation

Code for Figures: https://github.com/DCAN-Labs/nibabies-paper-code

## Author Contributions

Mathias Goncalves: Software, Conceptualization, Methodology, Writing - Original Draft; Julia Moser: Conceptualization, Investigation, Methodology, Writing - Original Draft; Thomas J. Madison: Software, Methodology, Writing - Original Draft; rae McCollum: Visualization, Formal Analysis, Data Curation/Processing, Validation, Writing - Original Draft; Jacob T. Lundquist: Data Curation/Processing, Validation, Writing - Original Draft; Begim Fayzullobekova: Data Curation, Validation; Lidia Hadera: Data Curation, Validation; Han H. Pham: Data Curation, Validation; Lucille A. Moore: Project administration, Writing - Review & Editing; Audrey Houghton: Software; Greg Conan: Software; Martin A. Styner: Methodology; Dimitrios Alexopoulos: Methodology; Christopher D. Smyser: Writing - Review & Editing; Sally M. Stoyell : Data Curation, Validation; Sanju Koirala: Data Curation, Writing - Review & Editing; Steven M. Nelson: Writing - Review & Editing; Kimberly B. Weldon: Project administration; Erik Lee: Methodology, Validation; Robert J. M. Hermosillo: Writing - Review & Editing; Luca Vizioli: Writing - Review & Editing; Essa Yacoub: Writing - Review & Editing; Gaurav H. Patel: Methodology; Juan Sanchez: Methodology; Kenneth Wengler: Methodology; Taylor Salo: Software, Methodology; Theodore Satterthwaite: Writing - Review & Editing; Jed T. Elison: Writing - Review & Editing; Christopher J. Markiewicz: Software, Writing - Review & Editing; Russell A. Poldrack: Writing - Review & Editing; Eric Feczko: Conceptualization, Supervision, Writing - Review & Editing; Oscar Esteban: Conceptualization, Software, Writing - Review & Editing; Damien A. Fair: Conceptualization, Supervision, Writing - Review & Editing;

## Acknowledgements

The authors acknowledge the Minnesota Supercomputing Institute (MSI) at the University of Minnesota for providing resources that contributed to the research results reported within this paper. URL: http://www.msi.umn.edu. Figure 1 icons provided by FlatIcon: https://flaticon.com/

## References

1. Li, X. et al. Moving beyond processing- and analysis-related variation in resting-state functional brain imaging. Nat. Hum. Behav. 1–15 (2024) doi:10.1038/s41562-024-01942-4.

2. Strother, S. C. Evaluating fMRI preprocessing pipelines. IEEE Eng. Med. Biol. Mag. 25, 27– 41 (2006).

3. Poldrack, R. A. et al. Scanning the horizon: towards transparent and reproducible neuroimaging research. Nat. Rev. Neurosci. 18, 115–126 (2017).

4. Kiar, G. et al. Align with the NMIND consortium for better neuroimaging. Nat. Hum. Behav. 7, 1027–1028 (2023).

5. Esteban, O. Standardized Preprocessing in Neuroimaging: Enhancing Reliability and Reproducibility. in Methods for analyzing large neuroimaging datasets, Neuromethods series (eds. Whelan, R. & Lemaître, H.) (Humana Press, New York, NY, 2025).

6. Esteban, O. et al. fMRIPrep: a robust preprocessing pipeline for functional MRI. Nat. Methods 16, 111–116 (2019).

7. Dubois, J. et al. MRI of the Neonatal Brain: A Review of Methodological Challenges and Neuroscientific Advances. J. Magn. Reson. Imaging 53, 1318–1343 (2021).

8. Korom, M. et al. Dear reviewers: Responses to common reviewer critiques about infant neuroimaging studies. Dev. Cogn. Neurosci. 53, 101055 (2021).

9. Kostović, I., Sedmak, G. & Judaš, M. Neural histology and neurogenesis of the human fetal and infant brain. Neuroimage 188, 743–773 (2019).

10. Dubois, J. et al. The early development of brain white matter: a review of imaging studies in fetuses, newborns and infants. Neuroscience 276, 48–71 (2014).

11. Eyre, M. et al. The Developing Human Connectome Project: typical and disrupted perinatal functional connectivity. Brain 144, 2199–2213 (2021).

12. Howell, B. R. et al. The UNC/UMN Baby Connectome Project (BCP): An overview of the study design and protocol development. Neuroimage 185, 891–905 (2019).

13. Volkow, N. D., Gordon, J. A. & Freund, M. P. The Healthy Brain and Child Development Study-Shedding Light on Opioid Exposure, COVID-19, and Health Disparities. JAMA Psychiatry 78, 471–472 (2021).

14. Esteban, O. et al. NiPreps: enabling the division of labor in neuroimaging beyond fMRIPrep. (2019).

15. Makropoulos, A. et al. The developing human connectome project: A minimal processing pipeline for neonatal cortical surface reconstruction. Neuroimage 173, 88–112 (2018).

16. Sturgeon, D. et al. DCAN-Labs Infant-Abcd-Bids-Pipeline. (2023). doi:10.5281/zenodo.7683282.

17. Sylvester, C. M. et al. Network-specific selectivity of functional connections in the neonatal brain. Cereb. Cortex (2022) doi:10.1093/cercor/bhac202.

18. Posse, S. Multi-echo acquisition. Neuroimage 62, 665–671 (2012).

19. Lynch, C. J. et al. Rapid Precision Functional Mapping of Individuals Using Multi-Echo fMRI. Cell Rep. 33, 108540 (2020).

20. Moser, J. et al. Multi-echo acquisition and thermal denoising advances precision functional imaging. Imaging Neuroscience 3, imag_a_00426 (2024).

21. Feczko, E. et al. Adolescent Brain Cognitive Development (ABCD) Community MRI Collection and Utilities. bioRxiv 2021.07.09.451638 (2021) doi:10.1101/2021.07.09.451638.

22. Sturgeon, D. et al. DCAN-Labs/abcd-Hcp-Pipeline. (2023). doi:10.5281/zenodo.10293494.

23. Glasser, M. F. et al. The minimal preprocessing pipelines for the Human Connectome Project. Neuroimage 80, 105–124 (2013).

24. Zhang, Y., Brady, M. & Smith, S. Segmentation of brain MR images through a hidden Markov random field model and the expectation-maximization algorithm. IEEE Trans. Med. Imaging 20, 45–57 (2001).

25. Wang, H. & Yushkevich, P. A. Multi-atlas segmentation with joint label fusion and corrective learning-an open source implementation. Front. Neuroinform. 7, 27 (2013).

26. Feczko, E. et al. Baby Open Brains: An open-source repository of infant brain segmentations. bioRxivorg 2024.10.02.616147 (2024) doi:10.1101/2024.10.02.616147.

27. Hendrickson, T. J. et al. BIBSNet: A deep learning Baby image brain segmentation network for MRI scans. bioRxivorg 2023.03.22.533696 (2024) doi:10.1101/2023.03.22.533696.

28. Fonov, V. S., Evans, A. C., McKinstry, R. C., Almli, C. R. & Collins, D. L. Unbiased nonlinear average age-appropriate brain templates from birth to adulthood. Neuroimage 47, S102 (2009).

29. Adamson, C. L. et al. Parcellation of the neonatal cortex using Surface-based Melbourne Children’s Regional Infant Brain atlases (M-CRIB-S). Sci. Rep. 10, 4359 (2020).

30. Nielsen, A. N. et al. Maturation of large-scale brain systems over the first month of life. Cereb. Cortex (2022) doi:10.1093/cercor/bhac242.

31. Zöllei, L., Iglesias, J. E., Ou, Y., Grant, P. E. & Fischl, B. Infant FreeSurfer: An automated segmentation and surface extraction pipeline for T1-weighted neuroimaging data of infants 0–2 years. Neuroimage 218, 116946 (2020).

32. Dale, A. M., Fischl, B. & Sereno, M. I. Cortical surface-based analysis. I. Segmentation and surface reconstruction. Neuroimage 9, 179–194 (1999).

33. Fischl, B. et al. Automatically parcellating the human cerebral cortex. Cereb. Cortex 14, 11– 22 (2004).

34. Markiewicz, C. J. et al. fmriprep-next: Preprocessing as a fit-transform model. in OHBM 2024 (2024).

35. DuPre, E. et al. TE-dependent analysis of multi-echo fMRI with tedana. J. Open Source Softw. 6, 3669 (2021).

36. Andersson, J. L. R., Skare, S. & Ashburner, J. How to correct susceptibility distortions in spin-echo echo-planar images: application to diffusion tensor imaging. Neuroimage 20, 870–888 (2003).

37. Goncalves, M. et al. NiTransforms: A Python tool to read, represent, manipulate, and apply dimensional spatial transforms. J. Open Source Softw. 6, 3459 (2021).

38. Glasser, M. F. et al. A multi-modal parcellation of human cerebral cortex. Nature 536, 171– 178 (2016).

39. Robinson, E. C. et al. MSM: a new flexible framework for Multimodal Surface Matching. Neuroimage 100, 414–426 (2014).

40. Robinson, E. C. et al. Multimodal surface matching with higher-order smoothness constraints. Neuroimage 167, 453–465 (2018).

41. Glasser, M. F. et al. The Human Connectome Project’s neuroimaging approach. Nat. Neurosci. 19, 1175–1187 (2016).

42. Mehta, K. et al. XCP-D: A Robust Pipeline for the post-processing of fMRI data. bioRxiv 2023.11.20.567926 (2023) doi:10.1101/2023.11.20.567926.

43. Bethlehem, R. A. I. et al. Brain charts for the human lifespan. Nature (2022) doi:10.1038/s41586-022-04554-y.

44. MacNicol, E. et al. Atlas-based brain extraction is robust across RAT MRI studies. in 2021 IEEE 18th International Symposium on Biomedical Imaging (ISBI) 312–315 (IEEE, 2021). doi:10.1109/isbi48211.2021.9433884.

45. Bridgen, P. et al. High resolution and contrast 7 tesla MR brain imaging of the neonate. Front. Radiol. 3, (2024).

46. Markiewicz, C. J. et al. fMRIPrep: A Robust Preprocessing Pipeline for Functional MRI. (Zenodo, 2024). doi:10.5281/ZENODO.12774221.

47. Gorgolewski, K. et al. Nipype: a flexible, lightweight and extensible neuroimaging data processing framework in python. Front. Neuroinform. 5, 13 (2011).

48. Abraham, A. et al. Machine learning for neuroimaging with scikit-learn. Front. Neuroinform. 8, 14 (2014).

49. Avants, B. B. et al. A reproducible evaluation of ANTs similarity metric performance in brain image registration. Neuroimage 54, 2033–2044 (2011).

50. Shi, F. et al. Infant brain atlases from neonates to 1- and 2-year-olds. PLoS One 6, e18746 (2011).

51. Chung, B. S. & Park, J. S. Automatic segmentation of true color sectioned images using FMRIB Software Library: First trial in brain, gray matter, and white matter. Clin. Anat. 33, 1197–1203 (2020).

52. Ciric, R. et al. TemplateFlow: FAIR-sharing of multi-scale, multi-species brain models. Nat. Methods 19, 1568–1571 (2022).

53. Jenkinson, M., Beckmann, C. F., Behrens, T. E. J., Woolrich, M. W. & Smith, S. M. FSL. Neuroimage 62, 782–790 (2012).

54. Hutton, C. et al. Image distortion correction in fMRI: A quantitative evaluation. Neuroimage 16, 217–240 (2002).

55. Greve, D. N. & Fischl, B. Accurate and robust brain image alignment using boundary-based registration. Neuroimage 48, 63–72 (2009).

56. Power, J. D. et al. Methods to detect, characterize, and remove motion artifact in resting state fMRI. Neuroimage 84, 320–341 (2014).

57. Jenkinson, M., Bannister, P., Brady, M. & Smith, S. Improved optimization for the robust and accurate linear registration and motion correction of brain images. Neuroimage 17, 825–841 (2002).

58. Behzadi, Y., Restom, K., Liau, J. & Liu, T. T. A component based noise correction method (CompCor) for BOLD and perfusion based fMRI. Neuroimage 37, 90–101 (2007).

59. Satterthwaite, T. D. et al. An improved framework for confound regression and filtering for control of motion artifact in the preprocessing of resting-state functional connectivity data. Neuroimage 64, 240–256 (2013).

60. Patriat, R., Reynolds, R. C. & Birn, R. M. An improved model of motion-related signal changes in fMRI. Neuroimage 144, 74–82 (2017).

61. Esteban, O., Goncalves, M., Markiewicz, C. J., Ghosh, S. S. & Poldrack, R. A. Software tool to read, represent, manipulate, and apply N-dimensional spatial transforms. in 2020 IEEE 17th International Symposium on Biomedical Imaging (ISBI) 709–712 (IEEE, 2020). doi:10.1109/isbi45749.2020.9098466.

62. Kundu, P., Inati, S. J., Evans, J. W., Luh, W.-M. & Bandettini, P. A. Differentiating BOLD and non-BOLD signals in fMRI time series using multi-echo EPI. Neuroimage 60, 1759– 1770 (2012).

63. Treiber, J. M. et al. Characterization and correction of geometric distortions in 814 diffusion weighted images. PLoS One 11, e0152472 (2016).

64. Fischl, B., Sereno, M. I. & Dale, A. M. Cortical surface-based analysis. II: Inflation, flattening, and a surface-based coordinate system. Neuroimage 9, 195–207 (1999).

65. Mugler, J. P., 3rd et al. Optimized single-slab three-dimensional spin-echo MR imaging of the brain. Radiology 216, 891–899 (2000).

66. Casey, B. J. et al. The Adolescent Brain Cognitive Development (ABCD) study: Imaging acquisition across 21 sites. Dev. Cogn. Neurosci. 32, 43–54 (2018).

67. Gordon, E. M. et al. Generation and Evaluation of a Cortical Area Parcellation from Resting-State Correlations. Cereb. Cortex 26, 288–303 (2016).

68. Demidenko, M., Mumford, J. & Poldrack, R. PyReliMRI: An Open-Source Python Tool for Estimates of Reliability in MRI Data. (Zenodo, 2024). doi:10.5281/ZENODO.12522260.

