## Supplemental Tables & Figures for "fMRIPrep Lifespan: Extending A Robust Pipeline for Functional MRI Preprocessing to Developmental Neuroimaging"

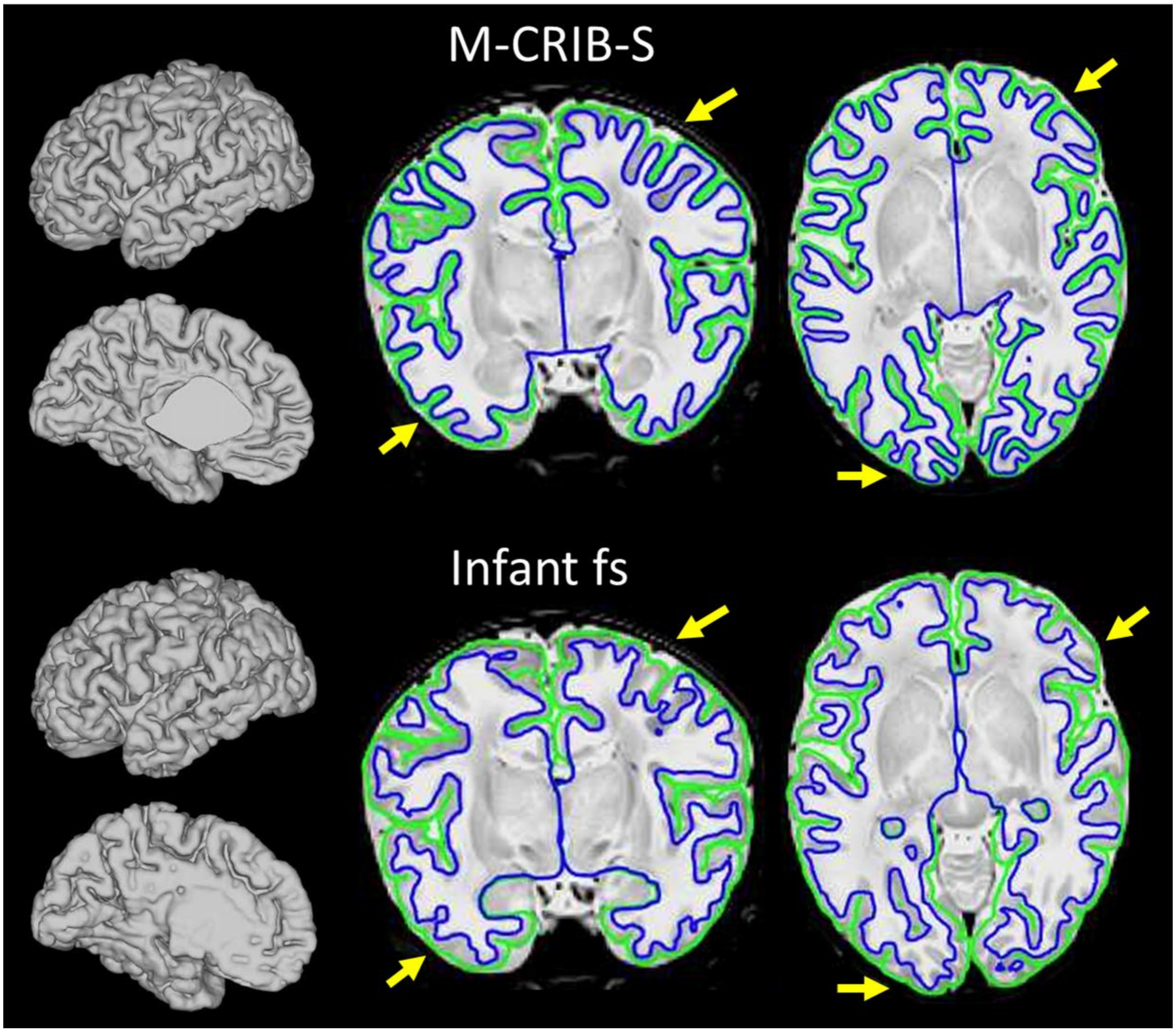


*Figure S1: Example of surface reconstruction performed with M-CRIB-S and Infant fs in a one month old subject. Both methods used the same segmentation generated by BIBSNet [27] as a basis. White matter surface shows more detail with M-CRIB-S while Infant fs misses the delineation of some smaller gyri (see arrows). M-CRIB-S is optimized for T2 based surface reconstruction at very young ages.*

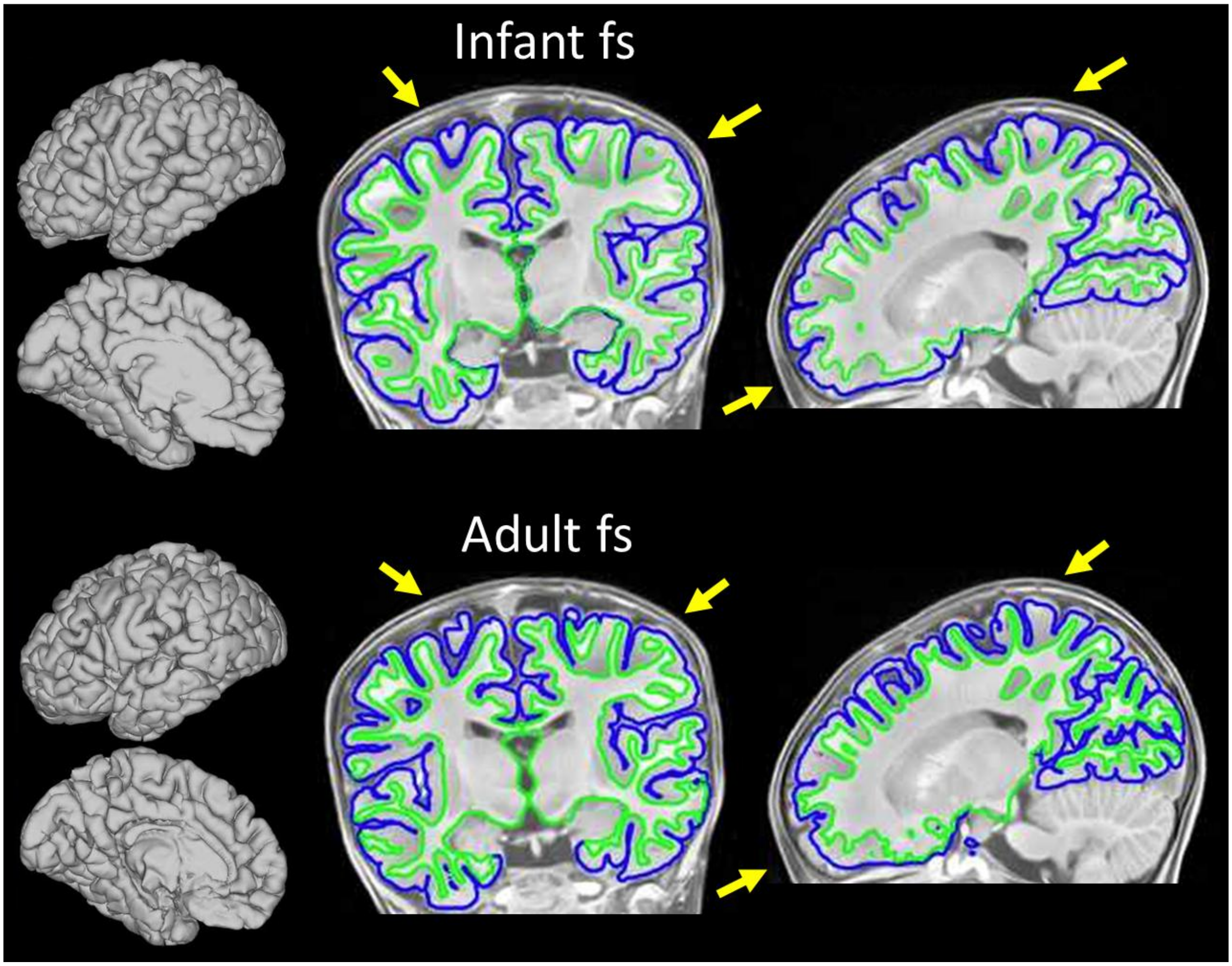


*Figure S2: Example of surface reconstruction performed with Infant fs and adult Freesurfer recon-all in a 17 month old subject. Infant fs uses the segmentation and brain mask generated by BIBSNet [27] as a basis. White matter surface shows more detail in some areas with Freesurfer recon-all however in this example, the Freesurfer based brain masking leads to gray matter cutoffs, which makes the surface created by Infant fs preferable.*

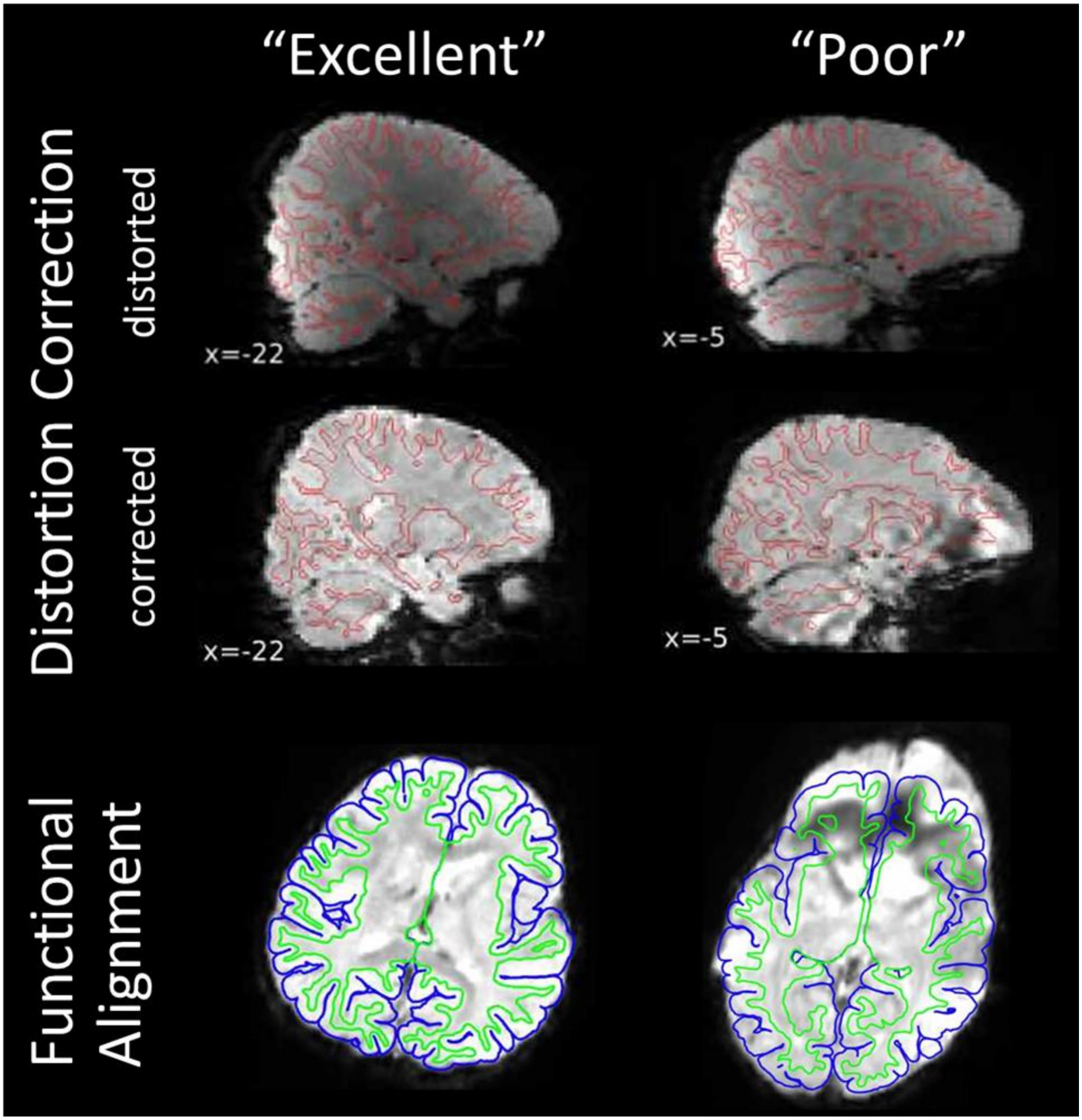


*Figure S3: Example derivatives rated as “excellent” and “poor” regarding distortion correction (top) and functional alignment (bottom). Examples are a 26 month old (left) and 20 month old (right). Data quality issues in the example on the right are caused by significant motion between fieldmaps.*

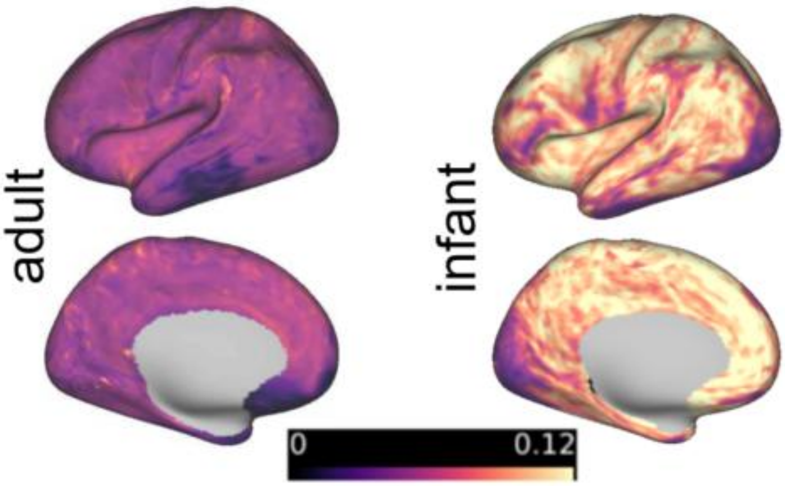


*Figure S4: Example for T2* maps (modified after [20]). Maps are projected onto the cortical surface, values represent T2* relaxation time in seconds.*

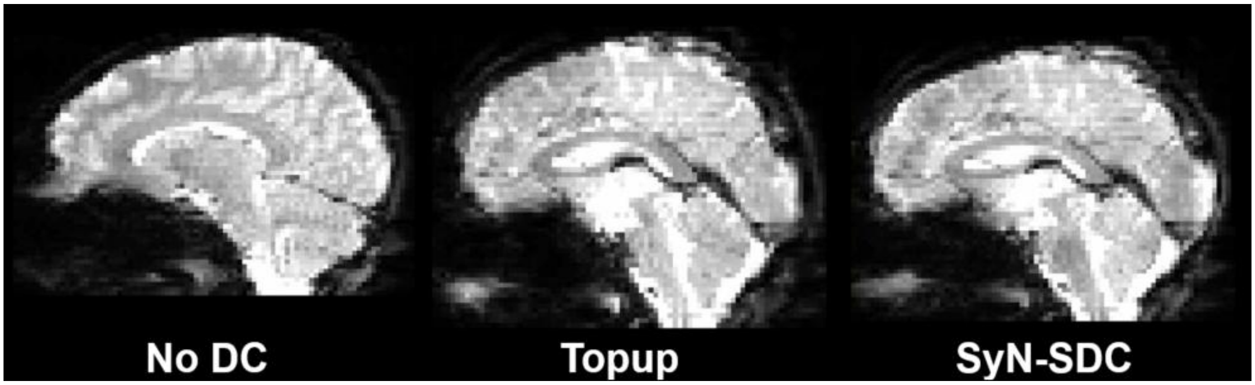


*Figure S5: A) Distortion correction (DC) in example subject, comparing no DC to FSL topup and fieldmapless SyN-SDC.*

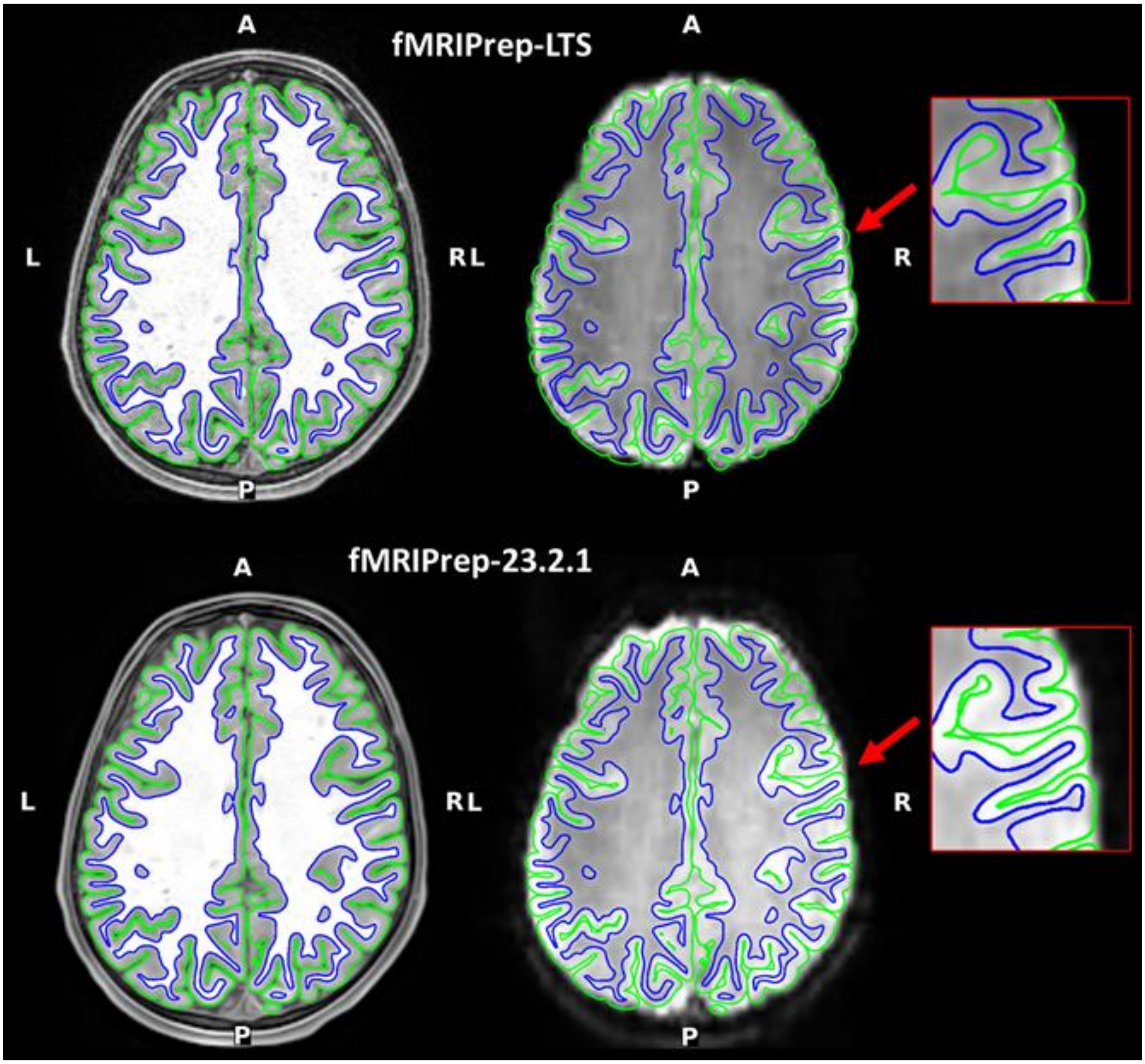

*Figure S6: Comparison of registration of surface to T1w image (left) and surface to BOLD image (right) between fMRIPrep-LTS and fMRIPrep-23.2.1 for a single ABCD resting-state run (5 minutes). Changes implemented between LTS and 23.2.1 (see Suppl. Table 1) improved BOLD registration.*

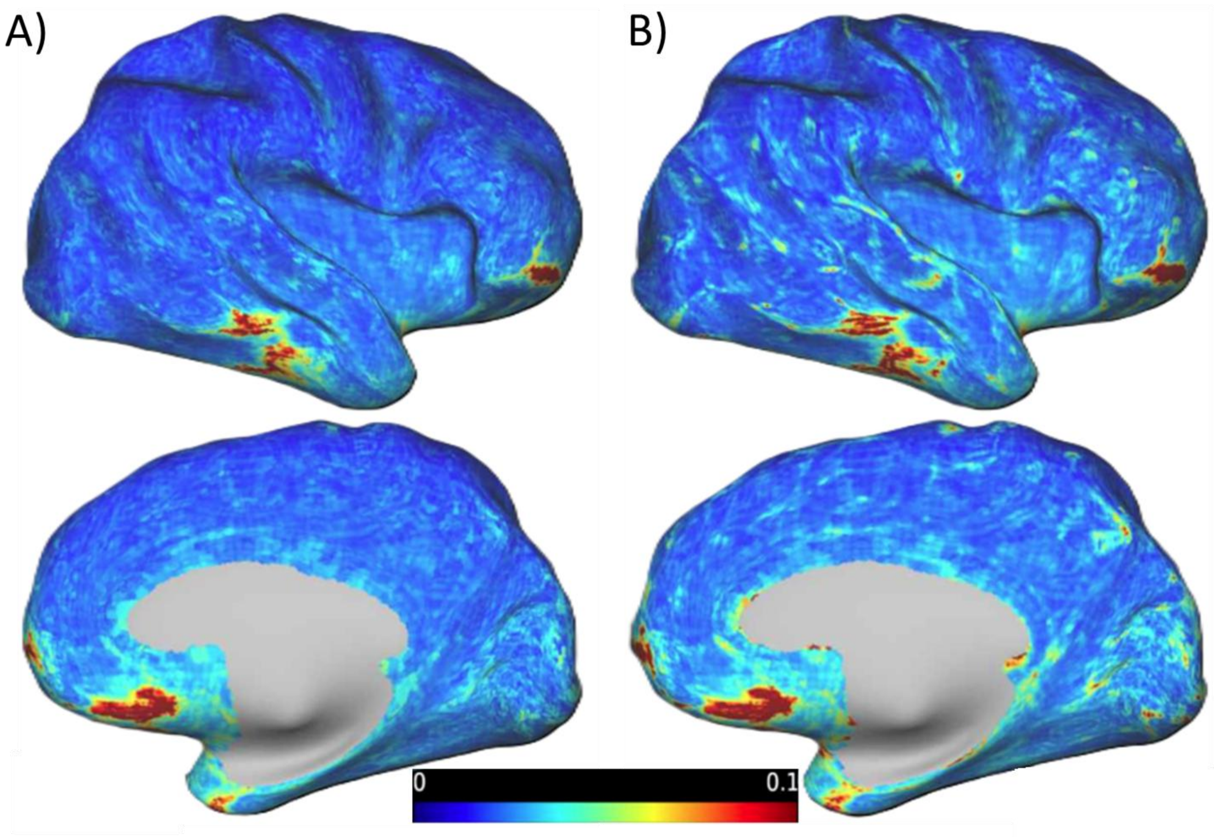


*Figure S7: Comparisons of outputs with (A) and without (B) 'good voxels' mask applied. Figure shows coefficient of variation (SD divided by mean) for a single ABCD resting-state run (5 minutes). The 'good voxels' mask excludes voxels with local peaks of temporal variation from the BOLD time-series*

| Participant ID | Session ID | Surface reconstruction Method | Surface recon-struction QC1 | Surface recon-struction QC2 | Spatial normalization QC1 | Spatial normalization QC2 | Distortion correction QC1 | Distortion correction QC2 | Functional alignment QC1 | Functional alignment QC2 |
|---|---|---|---|---|---|---|---|---|---|---|
| 421109 | 1mo | mcribs | 1 | 1 | 1 | 1 | 3 | 2 | 2 | 2 |
| 584381 | 1mo | mcribs | 1 | 1 | 1 | 1 | 1 | 1 | 1 | 1 |
| 960758 | 2mo | mcribs | 1 | 1 | 1 | 1 | 1 | 1 | 1 | 1 |
| 309615 | 3mo | mcribs | 1 | 1 | 1 | 1 | 1 | 1 | 1 | 1 |
| 375518 | 4mo | infantfs | 2 | 2 | 1 | 1 | 1 | 1 | 2 | 1 |
| 229768 | 5mo | infantfs | 1.5 | 1 | 1 | 1 | 1 | 1 | 1 | 1 |
| 197622 | 6mo | infantfs | 1 | 1.5 | 1 | 1 | 3 | 2 | 1.5 | 2 |
| 229768 | 8mo | infantfs | 1 | 1.5 | 1 | 1 | 1 | 1 | 1.5 | 1 |
| 132476 | 6mo | infantfs | 1 | 1 | 1 | 1 | 1 | 1 | 1.5 | 2 |
| 381606 | 5mo | infantfs | 1 | 1.5 | 1.5 | 1 | 1 | 1 | 1 | 1 |
| 116845 | 9mo | infantfs | 1 | 1.5 | 1 | 1 | 2 | 1 | 1.5 | 1 |
| 381606 | 11mo | infantfs | 2 | 1 | 1 | 1 | 1 | 1 | 1 | 1 |
| 530066 | 13mo | infantfs | 1 | 2 | 1 | 1 | 1 | 1 | 1 | 1 |
| 132476 | 15mo | adultfs | 3 | 1 | 2 | 1 | 1 | 1 | 1 | 1 |
| 100619 | 17mo | infantfs | 1 | 1.5 | 1 | 1 | 1 | 1 | 1 | 1 |
| 107842 | 19mo | infantfs | 1 | 1.5 | 1 | 1 | 1 | 1 | 1 | 1 |
| 200474 | 21mo | infantfs | 1 | 1.5 | 1 | 1 | 1 | 1 | 1 | 1 |
| 261266 | 23mo | infantfs | 1 | 1 | 1 | 1 | 1 | 1 | 1 | 1 |
| 105040 | 24mo | infantfs | 1 | 1.5 | 1 | 1 | 1 | 1 | 1 | 1 |
| 176851 | 20mo | infants | 1 | 1.5 | 1 | 1 | 2 | 2.5 | 2 | 2 |
| 353374 | 25mo | adultfs | 1 | 2 | 1 | 1 | 1 | 1 | 2 | 1 |
| 185373 | 26mo | infantfs | 1 | 1.5 | 1 | 1 | 1 | 1 | 1 | 1 |
| 505525 | 26mo | infantfs | 1 | 1.5 | 1 | 1 | 1 | 2 | 1.5 | 2 |
| 266394 | 27mo | infantfs | 1 | 1.5 | 1 | 1 | 1.5 | 2 | 1.5 | 2 |
| 764612 | 28mo | infantfs | 1 | 1.5 | 1 | 1 | 2 | 2.5 | 1.5 | 2.5 |
| 148796 | 29mo | infantfs | 1 | 1.5 | 1 | 1 | 1 | 1 | 1 | 1 |
| 418793 | 32mo | infantfs | 1 | 1 | 1 | 1 | 1 | 1 | 1 | 1 |
| 294064 | 34mo | infantfs | 1 | 1.5 | 1 | 1 | 1.5 | 1 | 1.5 | 1 |

| | | | | | | | | | | |
|---|---|---|---|---|---|---|---|---|---|---|
| 353374 | 37mo | infantfs | 1 | 1.5 | 1 | 1 | 1 | 1 | 1 | 1 |
| 676274 | 43mo | infantfs | 1 | 1 | 1 | 1 | 1 | 1 | 1 | 1 |

*Supplementary Table 1: Quality control outcomes for all participants separated into three age bins. Results were generated by six trained raters, each output was reviewed by two independent raters. 1= rated as high quality (“excellent”); 2=rated as usable with room for improvement (“acceptable”); 3= not of acceptable quality (“poor”);*

| **Version** | **Initial Release** | **Major Changes** |
|---|---|---|
| 23.2.x | January 2024 | Added MSMSulc-based registration to fsLR template; added option to ingress precomputed derivative files (instead of generating them as part of the fMRIPrep workflow)<br><br>SDCFlows updated to add Jacobian weighting step during unwarping, better masking of phase-difference and direct fieldmaps |
| 23.1.x | June 2023 | Overhaul of BOLD resampling workflows targeting fsLR spaces, based on HCP methods<br><br>CIFTI derivatives revised to use column-major ordering for subcortical grayordinates to match HCP 91k and 170k templates<br><br>Added T1w and T2w denoising step before surface reconstruction |
| 23.0.x | March 2023 | CIFTI dense scalar format output of FreeSurfer morphometric derivatives (curvature, cortical thickness, sulcal convexity) in fsLR space<br><br>Optional “goodvoxels” masking adapted from HCP methods<br><br>Addition of T2w volume derivatives in subject T1w space |
| 22.1.x | December 2022 | Head motion correction transforms and reference volumes added to functional derivatives<br><br>GIFTI format output of FreeSurfer morphometric derivatives<br><br>SDCFlows updated to 2.2.x, focusing on robustness of PEPolar workflow |
| 22.0.x | July 2022 | Move to FreeSurfer 7 (replacing 6.0.0) |

| | | |
|---|---|---|
| | | Estimated T2* maps and (in QA report) T2* histograms added to ME-EPI derivatives |
| 21.0.x | December 2021 | Adopt BIDS derivatives standard as the default output layout<br><br>SDCFlows major version update with new uniform API and adoption of FSL topup for PEPolar workflow (replacing AFNI 3dQwarp)<br><br>BOLD masking workflow revised to mitigate potential overcropping |

*Supplementary Table 2: Summary of the fMRIPrep change log from versions 21.0.x to 23.3.x*
